# Direction of ESCRT-III-dependent membrane bending emerges from bilayer asymmetry

**DOI:** 10.64898/2026.07.23.740341

**Authors:** Joshua Tran, Aurélien Roux

## Abstract

ESCRT-III is the cell’s principal machinery for reverse-topology membrane scission, cutting necks from within. Yet purified ESCRT-III predominantly assembles on the outside of membranes and bends them the opposite way. How one polymer chooses its direction is a long-standing paradox. Here, we show that ESCRT-III supplies the force, but bilayer asymmetry sets the direction. In budding yeast, sphingolipid asymmetry is required for efficient intraluminal vesicle formation: without it, cargo sorting is impaired and endosomal intermediates accumulate. In reconstituted membranes, imposing either lipid or protein asymmetry across the bilayer is sufficient to orient ESCRT-III-driven deformation. Leaflet-resolved fluorescence-lifetime imaging reveals that both act through a single mechanism: unequal lipid packing between leaflets generates differential tensile stress that dictates which way the membrane bends. Reconstituted membranes are typically symmetric; cellular membranes are not. This resolves the reverse-topology paradox and identifies bilayer asymmetry as a directional cue available to any membrane-remodeling machine.

## INTRODUCTION

Membrane remodeling is essential to cellular processes, from trafficking and signaling to cell division. While membrane budding and fission can occur in either direction relative to the cytosol, a distinct class of cellular processes requires membrane deformation away from the cytoplasm, into the lumen or extracellular space. The Endosomal Sorting Complex Required for Transport (ESCRT)-III machinery uniquely catalyzes reverse-topology membrane deformation and remains the only widely conserved system known to do so ^1^. Because fission changes membrane topology in the mathematical sense, the term “reverse-topology” can be ambiguous; we therefore refer to the direction of remodeling relative to the cytoplasm as “membrane orientation.”

Biochemical, structural, and *in vitro* reconstitution studies have established that ESCRT-III components assemble into polymers and VPS4-driven ATP hydrolysis remodel and perform membrane scission ^2–8^. In cells, ESCRT-III is dynamically recruited with precise spatiotemporal control to drive diverse processes essential for homeostasis, including multivesicular body biogenesis ^9,10^, cytokinetic abscission ^11,12^, maintenance of lysosomal ^13,14^ and plasma membrane integrity ^15^, nuclear envelope reformation ^16,17^, and viral budding ^18–20^.

Yet a fundamental discrepancy persists. *In vitro*, ESCRT-III mediates bending and scission in both membrane orientations ^1,21–23^ as expected from its buckling mechanism ^22^. However, this flexibility is not unbiased as the polymer intrinsically favors outwards budding, and inward deformation emerges only when membranes are self-assembled onto preformed ESCRT-III polymers ^24,25^. In contrast, in cells, ESCRT-III-mediated bending is strikingly unidirectional, consistently directed away from the cytosol, the orientation the polymer disfavors *in vitro*. The discrepancy therefore does not lie in ESCRT-III itself, which is the same protein in both settings, but in the membrane it acts on. Something present in cellular membranes, and absent from reconstituted ones, must override the intrinsic preference of the polymer. What that is, and how it couples to polymer mechanics, remains unresolved.

Cellular membranes are lipid bilayers exhibiting pronounced asymmetry between their two leaflets: lipid and protein composition and, consequently, mechanical properties differ substantially between extracellular and cytosolic leaflets ^26–28^. By contrast, the model membranes used for in vitro reconstitution are typically symmetric. ESCRT-III proteins assemble and act exclusively from the cytosolic face of the membrane, yet drive vesicle formation away from it, into the lumen or extracellular space. This arrangement raises the possibility that pre-existing bilayer asymmetry contributes to the orientation of ESCRT-III-mediated membrane deformation. Indeed, asymmetric lipid organization can generate membrane curvature, as illustrated by Shiga toxin, which clusters lipids within one leaflet and drives bending toward the opposite side ^29^. Thus, differences in the molecular area and physical state of the two leaflets can provide an intrinsic bias for membrane curvature. We therefore hypothesized that ESCRT-III couples its polymer mechanics to this pre-existing bilayer asymmetry to set the direction of membrane bending and vesiculation.

## RESULTS

### Sphingolipid and phospholipid homeostasis regulates ILV formation in budding yeast

If bilayer asymmetry contributes to the physical environment in which ESCRT-III acts, then perturbing it in cells should affect ESCRT-dependent cargo sorting. We first tested this prediction using the internalization and vacuolar delivery of Mup1p in *Saccharomyces cerevisiae*. Upon methionine stimulation, Mup1p is endocytosed and sorted into intraluminal vesicles (ILVs) at the multivesicular body in an ESCRT-III-dependent manner ^30–33^. Defects in ILV formation prevent efficient delivery of Mup1p into the vacuolar lumen, resulting instead in its accumulation at the vacuolar limiting membrane and at stalled intermediate compartments (Figure 1A).

**Figure 1.**
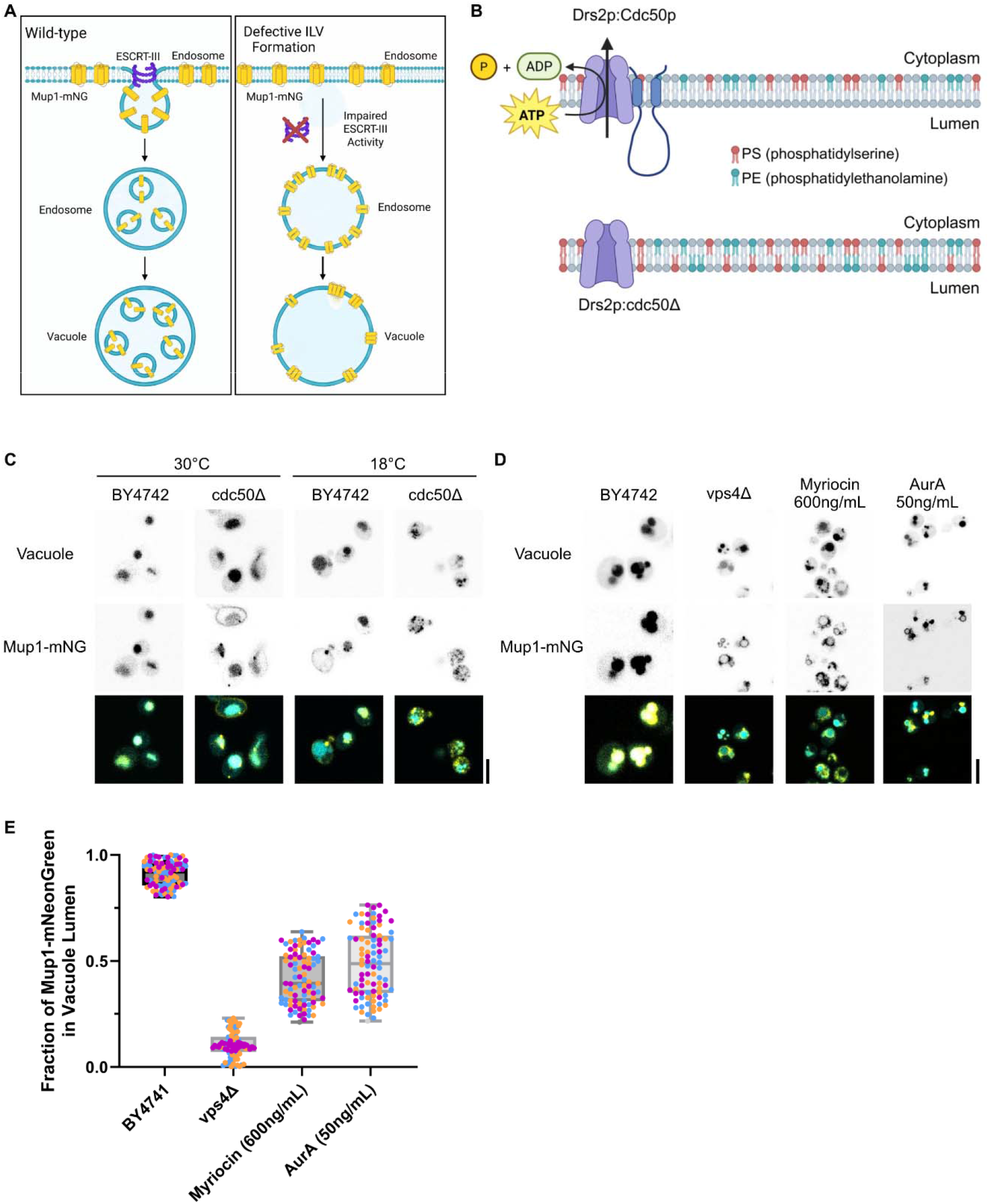
Sphingolipid and phospholipid homeostasis regulates ILV formation in budding yeast. **(A)** Schematic of ESCRT-III–dependent intraluminal vesicle (ILV) formation. **(B)** Schematic of phospholipid asymmetry generated by the P4-ATPase flippase complex Drs2p–Cdc50p. **(C)** Mup1p– mNeonGreen sorting after a methionine pulse in BY4742 cells and cdc50Δ mutants at 30°C (left) and 18°C (right). **(D)** Mup1p–mNeonGreen sorting after a methionine pulse in BY4742 cells treated with DMSO (vehicle), myriocin (600 ng/mL), or aureobasidin A (50 ng/mL), and in vps4Δ mutants. In (C) and (D), cargo is shown in yellow and the vacuole lumen in cyan. **(E)** Fraction of Mup1p–mNeonGreen signal in the vacuole (vacuolar/total cell signal) for the conditions in (D). Each dot represents one cell (30 cells per experiment); colors denote three independent experiments. Box plots show the median (center line), interquartile range (box limits), and minimum to maximum (whiskers). Scale bars, 2 µm. Schematics created with BioRender.com.

To perturb phospholipid asymmetry, we analyzed endocytic trafficking and degradation of Mup1p– mNeonGreen in *cdc50*Δ mutants. Cdc50p, in complex with the P4-ATPase Drs2p, forms a flippase that maintains endosomal phospholipid asymmetry by catalyzing the inward translocation of phosphatidylserine (PS) and phosphatidylethanolamine (PE) to the cytosolic leaflet ^34–36^ (Figure 1B).

Because PE has a relatively small headgroup, its enrichment in the cytosolic leaflet can contribute to leaflet area asymmetry and thereby influence membrane curvature. We therefore asked whether disrupting this lipid asymmetry alters the efficiency of ESCRT-dependent ILV formation. To trigger Mup1p internalization, cells were grown in methionine-deficient medium and acutely stimulated with 20 µg/mL methionine. In wild-type cells, Mup1p–mNG was efficiently delivered to the 7-amino-4-chloromethylcoumarin (CMAC)-labeled vacuolar lumen. In contrast, in *cdc50*Δ mutants, vacuolar delivery remained detectable, but small cytoplasmic Mup1p-positive puncta accumulated and persisted throughout the imaging period (Figure 1C). Notably, clear vacuolar localization of cargo was observed in some cells. Thus, loss of Cdc50p did not abolish Mup1p trafficking to the vacuole but instead produced a partial trafficking defect characterized by persistent cargo-positive intermediates, consistent with inefficient ILV formation.

Previous work has reported that *cdc50*Δ mutants exhibit strong cold sensitivity, along with delayed ESCRT-dependent degradation of cargoes such as Ste3p ^34^. We therefore examined Mup1p– mNeonGreen trafficking at 18°C to determine whether membrane asymmetry becomes more important under reduced temperature conditions. At 18°C, wild-type cells still efficiently delivered Mup1p– mNeonGreen to the vacuolar lumen, but with slower kinetics than at 30°C. In contrast, *cdc50*Δ mutants displayed a pronounced trafficking defect, with Mup1p–mNeonGreen largely excluded from the vacuolar lumen and accumulating in numerous intracellular puncta throughout the cell (Figure 1C).

This cold-sensitive phenotype suggested that perturbing phospholipid asymmetry makes endosomal cargo delivery particularly sensitive to changes in membrane physical state. Because temperature changes membrane properties, including lipid packing and molecular order, we reasoned that the pronounced defect in *cdc50*Δ mutants at 18°C could reflect an increased dependence of endosomal sorting on membrane organization. We therefore asked whether sphingolipid homeostasis, which strongly influences membrane organization and physical properties ^37^, contributes to this temperature-dependent trafficking defect.

To perturb the asymmetric sphingolipid pool, we treated wild-type cells with myriocin, an inhibitor of serine palmitoyltransferase (SPT), which blocks the first committed step in sphingolipid synthesis (Figure S1A). Complex sphingolipids are synthesized on the luminal side of the Golgi and subsequently become enriched in the exoplasmic/luminal leaflet of the plasma membrane and endosomal membranes, thereby contributing to bilayer asymmetry. Compared to DMSO-treated controls, myriocin markedly reduced cargo delivery into the vacuolar lumen and caused cargo-positive puncta to accumulate adjacent to the vacuole (Figure 1D). Unlike *vps4*Δ mutants, which exhibit a near-complete block in cargo delivery, myriocin-treated cells retained partial sorting capacity, indicating that depletion of sphingolipids reduces the efficiency of ESCRT-III-dependent ILV formation (Figure 1E). We further inhibited complex sphingolipid synthesis with aureobasidin A (AbA), which blocks ceramide incorporation into downstream sphingolipids, and observed a comparable reduction in cargo delivery (Figure 1E, F). Consistent with impaired ILV-mediated degradation, myriocin also increased canavanine sensitivity, as both WT and *cdc50*Δ mutants showed markedly reduced growth in the presence of canavanine following myriocin treatment (Figure S1B). Together, these perturbations indicate that sphingolipid homeostasis contributes to the efficiency of ESCRT-dependent ILV formation.

Taken together, these experiments show that ESCRT-dependent Mup1p trafficking remains functional under basal conditions but becomes strongly sensitive to perturbations of phospholipid and sphingolipid homeostasis, particularly when membrane physical state is altered by low temperature. The partial retention of cargo delivery after sphingolipid depletion distinguishes these phenotypes from the near-complete trafficking block caused by loss of Vps4 and indicates that altered lipid organization does not simply abolish ESCRT function. These perturbations establish that ESCRT-dependent ILV formation is sensitive to bilayer lipid composition and asymmetry. They cannot, however, reveal whether asymmetry sets the direction of bending, because every ILV in these cells buds in the same direction: there is no condition in which the sign of the deformation can be read out. Testing directionality requires a system in which the sign of the asymmetry can be imposed at will. We therefore turned to *in vitro* reconstitution.

### Asymmetric sphingolipid distribution drives inwards invagination in vitro

To determine whether lipid asymmetry is sufficient to bias ESCRT-III driven membrane deformation, we reconstituted Snf7p, the first polymerizing subunit of yeast ESCRT-III, activity on giant unilamellar vesicles (GUVs) composed of DOPC (44.95mol%), DOPS (40mol%), brain sphingomyelin (bSM) (15mol%), and the far-red lipid dye DiD (0.05mol%). Because this reconstitution contains Snf7p alone, without Vps4 or ATP, it reports on the bending step of the reaction rather than on scission. We therefore use the direction of membrane deformation, not fission, as our readout throughout. We perturbed sphingolipid organization in the GUV using bacterial sphingomyelinase (bSMase), which enzymatically converts sphingomyelin into ceramide (Figure 2A). GUVs were prepared with sphingomyelin in both leaflets. Because bSMase is added externally and does not cross the bilayer, hydrolysis is confined to the outer leaflet, generating a defined compositional asymmetry in a membrane that began symmetric. Topologically, the outer leaflet of the GUV corresponds to the luminal leaflet of the endosome, so inward deformation of the GUV corresponds to ILV-directed budding in cells. bSMase therefore imposes a leaflet-specific change in composition, molecular area, and preferred curvature, with a defined sign. Previous reports have suggested that active sphingomyelin conversion to ceramide can be a mechanism for membrane vesiculation independently of protein machineries ^37–39^.

**Figure 2.**
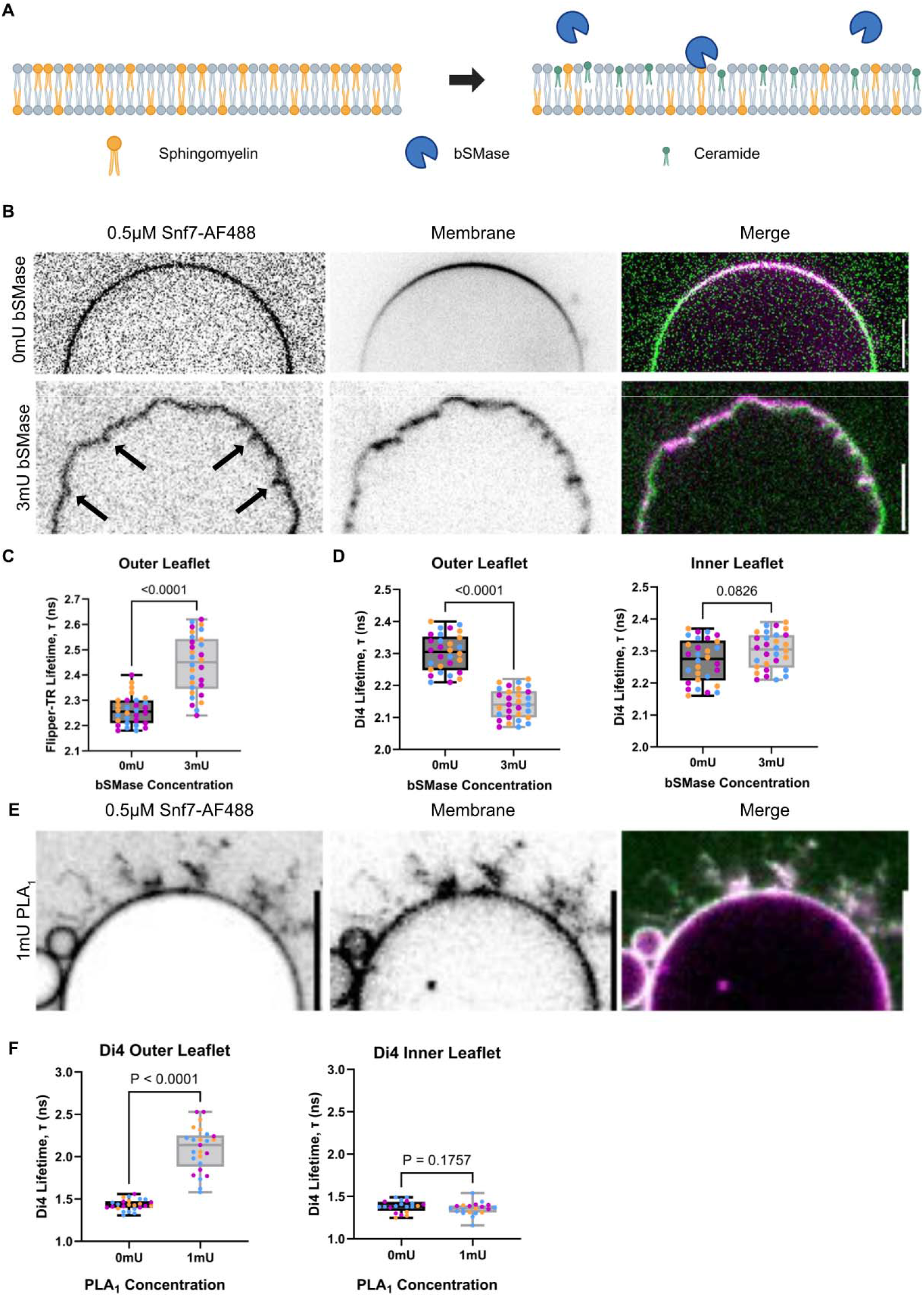
Asymmetric sphingolipid distribution drives inwards invagination *in vitro*. **(A)** Schematic of sphingomyelin conversion to ceramide by bSMase on giant unilamellar vesicles (GUVs). **(B)** Alexa Fluor 488–labeled Snf7p (0.5 µM) bound to GUVs composed of DOPC (44.95 mol%), DOPS (40 mol%), brain sphingomyelin (bSM, 15 mol%), and DiD (0.05 mol%), without (top row) or after (bottom row) bSMase treatment. **(C)** Fluorescence lifetime of Flipper-TR in the GUV outer leaflet before and after bSMase treatment (3 mU). **(D)** Fluorescence lifetime of Di-4 in the GUV outer and inner leaflets before and after bSMase treatment (3 mU). **(E)** Alexa Fluor 488–labeled Snf7p (0.5 µM) bound to GUVs as in (B) in the presence of PLA1 (1 mU). **(F)** Fluorescence lifetime of Di-4 in the GUV outer and inner leaflets before and after PLA1 treatment (1 mU). In (C), (D), and (F), each dot represents one GUV (10 GUVs per experiment); colors denote three independent experiments. Box plots show the median (center line), interquartile range (box limits), and minimum to maximum (whiskers). Lifetimes are in ns. Statistical significance was assessed by two-sided Mann–Whitney U test on GUVs pooled across experiments (n = 30 per condition); exact P values are shown on the plots. Scale bars, 5 µm. Schematic created with BioRender.com.

In the absence of enzymatic perturbation, fluorescently labeled Snf7p (Snf7–AF488) bound to GUV membranes but did not induce detectable out-of-plane deformation, consistent with previous reports of Snf7p polymerizing into predominantly flat spirals on membranes (Figure 2B, top row) ^22,40,41^. To test the response of ESCRT-III to enzymatically treated membranes, we used a concentration of bSMase that does not drive detectable spontaneous budding (Figure S2C). Although these GUVs remained macroscopically undeformed, sphingomyelin hydrolysis selectively altered the composition and physical state of the enzyme-exposed leaflet, establishing an asymmetric preferred area and, consequently, a spontaneous-curvature bias. We therefore reasoned that the absence of detectable deformation represents a pseudo-steady state in which the membrane asymmetry is established but does not, by itself, produce macroscopic curvature, consistent with work done with methyl-β-cyclodextrin ^42^. Upon addition of Snf7-AF488 to bSMase-treated GUVs, we observed robust formation of inward membrane invaginations (Figure 2B, bottom row, arrows). No comparable deformation occurred with bSMase alone under the same conditions or with Snf7p in the absence of bSMase. Thus, the emergence of inward deformation upon Snf7p assembly provides a morphological readout of the asymmetric membrane state generated by sphingomyelin hydrolysis and demonstrates that this pre-existing curvature bias directs ESCRT-III-mediated remodeling toward inward bending.

To determine whether ceramide itself is sufficient to reproduce this effect, independent of its asymmetric distribution between the two leaflets, we generated GUVs containing symmetric C18 ceramide (Figure S2D). Despite the presence of a lipid species with a strong intrinsic curvature preference, Snf7p failed to induce detectable membrane deformation in these symmetric membranes. Thus, symmetric ceramide did not reproduce the directional ESCRT response under these conditions.

Preferred direction of bending can be theoretically driven by a stress asymmetry between the two leaflets ^28,43,44^. We thus next asked whether enzymatic sphingomyelin remodeling generates a stress asymmetry between the two leaflets of the membrane. To resolve leaflet-specific changes in membrane physical state, we employed fluorescence lifetime imaging microscopy (FLIM) using two complementary fluorescent probes.

We first used Flipper-TR, a mechanosensitive probe whose fluorescence lifetime (τ) responds to changes in the lateral mechanical state of the membrane ^45^. In single-phase model membranes, increased lateral compression of the membrane is associated with increased Flipper-TR fluorescence lifetime, reflecting planarization of the probe ^46,47^. Following addition of bSMase to GUVs, we observed an increase in Flipper-TR lifetime in the outer leaflet. This response is consistent with increased lateral compression of the outer leaflet resulting from the conversion of sphingomyelin to the smaller ceramide molecule (Figure 2C). To evaluate the inner leaflet, however, Flipper-TR turned out to be difficult to extract from the membrane.

We therefore turned to Di-4-ANEPPDHQ (Di4), a membrane-packing-sensitive fluorescent probe that localizes near the membrane–water interface and whose fluorescence lifetime (τ) increases with lipid packing ^27,48,49^. Di4 was retained in the membrane while preserving leaflet-specific labeling (Figure S2A), enabling us to assess the physical state of each leaflet. Following bSMase treatment, Di4 lifetime decreased in the external leaflet, consistent with reduced molecular order at the membrane–water interface following sphingomyelin hydrolysis (Figure 2D, left). By contrast, after selectively removing the outer-leaflet probe with fatty-acid-free BSA, Di4 lifetime in the inner leaflet remained unchanged (Figure 2D, right). Thus, sphingomyelin hydrolysis selectively altered the composition and interfacial lipid packing of the enzyme-exposed leaflet, potentially generating a mechanical asymmetry between the two leaflets.

These changes can be understood from the molecular consequences of sphingomyelin hydrolysis. Cleavage of the phosphocholine headgroup converts sphingomyelin, with a molecular area of approximately 51 Å² ^27^, into ceramide, with an effective molecular area of approximately 35 Å² ^50^. This conversion reduces the preferred molecular area of the outer leaflet while altering its interfacial lipid organization. Because hydrolysis occurs selectively in the outer leaflet, the resulting area mismatch between the leaflets generates differential stress through bilayer coupling, establishing distinct mechanical states that bias spontaneous curvature even in the absence of detectable macroscopic deformation, consistent with other work ^42^. Within the bilayer-couple framework, the reduced preferred area of the outer leaflet favors inward bending, with the larger-preferred-area inner leaflet occupying the convex surface. To test whether an opposing perturbation of membrane organization could reverse this directional bias, we used PLA1 (phospholipase A1), which hydrolyzes phosphatidylcholine (PC) to generate lysophospholipids and free fatty acids, and can tubulate membranes ^51^. PLA1 treatment induced outward tubulation (Figure 2E) and an opposite change in Di4-measured interfacial organization (Figure 2F), consistent with an opposing mechanical bias. Together, these findings support a model in which asymmetric lipid remodeling establishes differential leaflet stress that biases curvature direction, while ESCRT-III assembly drives the membrane deformation that expresses this bias.

### Asymmetric cargo crowding controls directional bending in vitro

During ILV formation, cargoes are recognized, clustered, and internalized by the ESCRT machinery. Cargoes have their steric volume asymmetrically distributed across the bilayer, and this may create a leaflet stress asymmetry as more crowding of cargoes may occur on one side of the membrane. We therefore asked whether cargo distribution across the membrane bilayer, and its resulting asymmetry, can encode membrane remodeling directionality (Figure 3A). To test this, we reconstituted fluorescent 6×His– tetraubiquitin (linear chain) constructs on giant unilamellar vesicles (GUVs) containing DGS-NTA(Ni) (Ni-NTA) lipids, hereafter referred to as His-Ub4 cargo and monitored resulting membrane deformations. This tetraubiquitin construct has previously been used to probe ESCRT-dependent cargo recognition and clustering ^52^.

**Figure 3.**
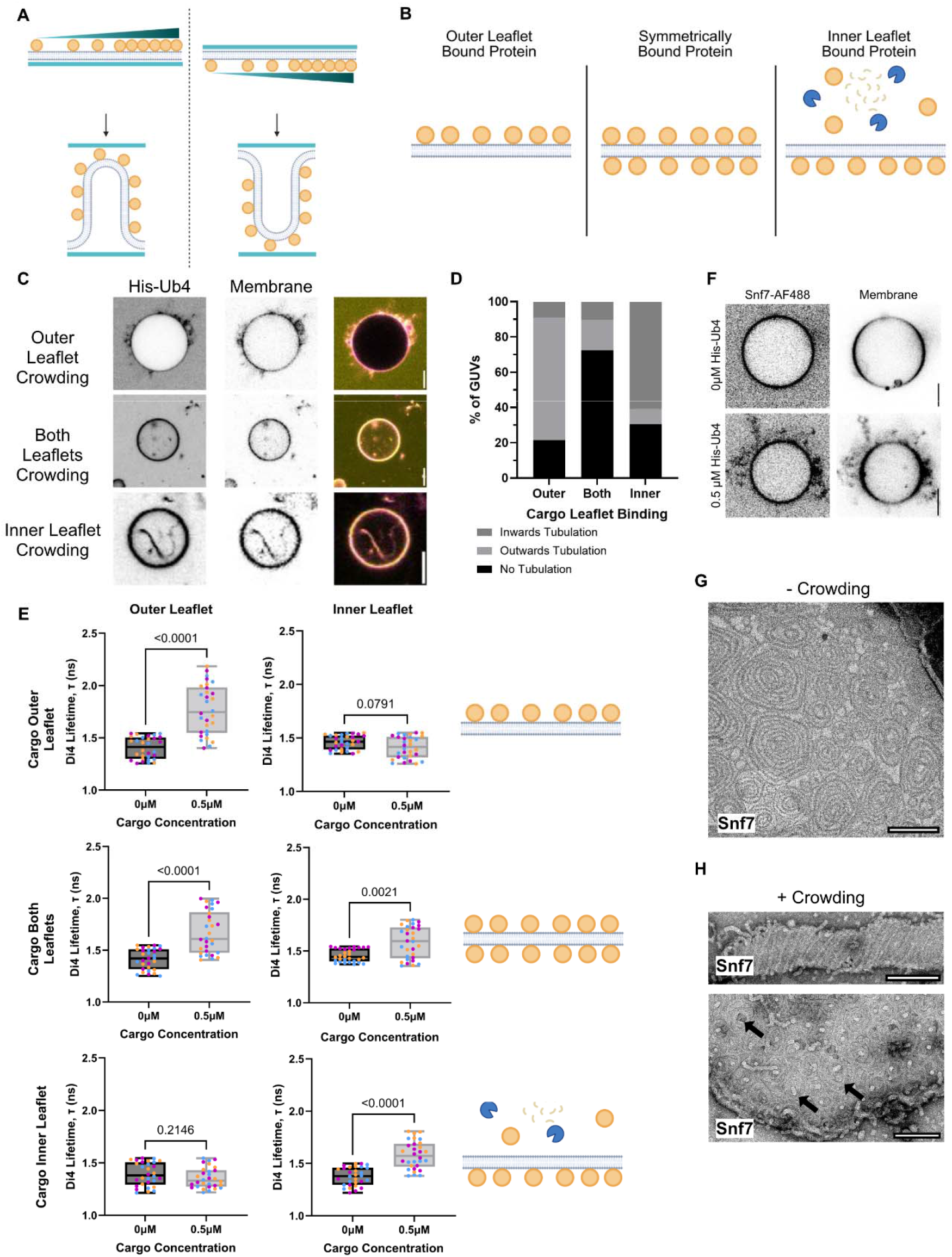
Asymmetric cargo crowding controls directional bending in vitro. **(A)** Schematic of asymmetric cargo crowding and directional bending. **(B)** Schematic of leaflet-specific control of His-Ub4 cargo crowding by external addition, symmetric addition, or proteinase K cleavage. **(C)** Representative images of asymmetric and symmetric cargo crowding on GUVs containing 11.5 mol% DGS-NTA(Ni). **(D)** Quantification of directional tubulation in (C). **(E)** Fluorescence lifetime of Di-4 in the GUV outer and inner leaflets with cargo bound to the outer leaflet (top row), both leaflets (middle row), or the inner leaflet (bottom row). **(F)** Representative images of Snf7p (0.5 µM, sub-tubulating concentration) bound to GUVs with or without cargo. **(G)** Representative negative-stain electron microscopy (EM) image of Snf7p spirals on flat membranes. **(H)** Representative negative-stain EM image of Snf7p bound to cargo-crowded membrane tubes and large unilamellar vesicle (LUV) membranes. In (E), each dot represents one GUV (10 GUVs per experiment); colors denote three independent experiments. Box plots show the median (center line), interquartile range (box limits), and minimum to maximum (whiskers). Lifetimes are in ns. Statistical significance was assessed by two-sided Mann–Whitney U test on GUVs pooled across experiments (n = 30 per condition); exact P values are shown on the plot. Scale bars, 5 µm (C and F) and 50 nm (G and H). Schematics created with BioRender.com.

Addition of 1 µM His-Ub4 to the outer leaflet of GUVs (11.5 mol% Ni-NTA, 40 mol% DOPS, 48.45 mol% DOPC, 0.05 mol% DiD) induced robust outward membrane tubulation, with curvature directed away from the cargo-enriched leaflet (Figure 3C, D), consistent with a crowding-induced lateral stress on the bound membrane leaflet ^53^. To test whether asymmetric cargo distribution is required for this effect, His-Ub4 was incorporated during GUV formation, allowing binding to both leaflets. Under these conditions, membrane tubulation was strongly reduced (Figure 3C, D), indicating that symmetric distribution of cargo across the bilayer suppresses curvature generation. We next asked if inward tubulation could be generated by asymmetric localization of the cargo to the inner leaflet of GUVs. For this, on GUV symmetrical coated with his-Ub4, externally bound His-Ub4 was selectively removed with proteinase K (final concentration 0.2 mg/mL as described in ^54^), restricting the remaining cargo to the inner leaflet. Moving the same molecule to the opposite leaflet inverted the direction of deformation: outer-leaflet cargo produced outward tubulation, inner-leaflet cargo produced inward tubulation, with no change in cargo identity, membrane composition, or buffer. The sign of the deformation therefore follows the sign of the asymmetry (Figure 3B–D).

We next asked how asymmetric protein binding alters the physical state of each membrane leaflet. We therefore used the same leaflet-specific FLIM measurements established above. Di4 fluorescence lifetime increased selectively in the cargo-bound leaflet, consistent with increased molecular order and packing at the membrane–water interface, whereas the opposing leaflet remained unchanged within detectable limits (Figure 3E). Independently, Flipper-TR fluorescence lifetime decreased in the cargo-bound outer leaflet, consistent with probe deplanarization in single-phase membranes and consistent with increased lateral expansion of the leaflet and increased lateral tension (Figure S3C). Thus, asymmetric protein crowding produces coordinated changes in the physical state of the cargo-bound leaflet, increasing molecular packing at the membrane–water interface while increasing its effective lateral area and producing a Flipper response consistent with increased lateral tension.

Together, these measurements indicate that asymmetric cargo crowding generates a mechanical asymmetry across the bilayer. The resulting difference in leaflet physical state provides a directional bias for membrane curvature, with deformation occurring away from the cargo-enriched leaflet.

We then aimed at testing whether this mechanical asymmetry generated by cargo crowding could direct ESCRT-III-mediated membrane deformation. For this, we added Snf7p to GUV coated with His-Ub4 in various ways. Importantly, membrane deformation was not observed when Snf7p was added alone at the tested concentrations, and we chose conditions of cargo binding at sub-tubulating levels, providing undetectable curvature in the absence of ESCRT-III (Figure 3G). When ESCRT-III polymerization and cargo crowding were combined on the same membrane, however, robust membrane bending was observed. When cargo was restricted to the outer leaflet, deformation occurred away from the cargo-enriched side, resulting in outward tubulation (Figure 3F, bottom). In the absence of protein crowding, Snf7p assembled into flat spirals on membranes without detectable out-of-plane deformation (Figure 3G). In contrast, in the presence of crowding cargoes, negative-stain electron microscopy revealed that Snf7p formed ring-like polymers tightly encircling tubular membranes, oriented perpendicular to the axis of elongation, consistent with ESCRT-III filaments directly contributing to membrane curvature generation (Figure 3H, top). Moreover, membrane protrusions extending outward from the centers of Snf7p spirals were observed on LUVs (Figure 3H, bottom). These structures closely resemble protrusions previously reported during VPS4-mediated turnover of ESCRT-III polymers, suggesting that protein crowding, when coupled to Snf7p polymerization, can similarly drive out-of-plane membrane deformation.

Together, these experiments establish that the direction of ESCRT-III-mediated membrane deformation depends on the leaflet in which protein crowding is imposed. Steric interactions between membrane-bound cargoes expand the effective area occupied by the cargo-bearing leaflet, producing a leaflet-specific change in membrane physical state consistent with increased lateral tension and increased molecular packing at the membrane–water interface. Within the bilayer-couple framework, this leaflet area mismatch generates a spontaneous-curvature bias in which the expanded leaflet preferentially occupies the convex surface.

Neither sub-tubulating cargo crowding nor Snf7p polymerization alone generated detectable macroscopic curvature, indicating that the asymmetric membrane state can be established without visible deformation. ESCRT-III polymerization therefore acts on a pre-existing curvature bias and supplies the additional mechanical work required to express that bias as macroscopic membrane deformation. These results show that membrane asymmetry is sufficient to bias the direction of active ESCRT-mediated remodeling and can be reversed by changing the leaflet distribution of the same molecular crowder.

### Asymmetric cargo topology affects cargo sorting efficiency in cellulo

We next wondered if the same mechanism of remodeling directionality was at play in vivo. To answer this, we tested whether asymmetric cargo crowding influences ILV formation in the budding yeast. To uncouple ubiquitin-dependent cargo recognition from domain topology, we used an Hse1-DUb strain in which the ESCRT-0 component Hse1 is fused with a deubiquitinase, providing deubiquination of cargoes gathered by ESCRT-0, allowing them to exit the ESCRT pathway. This strain shows a complete block of ILV formation despite functional ESCRT-III ^55^. ILV formation and cargo incorporation can be restored by fusing a non-cleavable ubiquitin directly to the cargo ^55^ (Figure 4A, B). We therefore engineered a set of synthetic cargoes based on the single-pass transmembrane domain of Cps1p, each carrying an in-frame ubiquitin on the cytosolic domain to maintain ESCRT recognition. Two copies of mouse metallothionein increased the bulk of the cargo domain and provided a defined protein mass, while mScarlet-i3 enabled live-cell imaging (Figure S4A). We varied the distribution of this protein bulk across the membrane to generate cytosolic-large, luminal-large, and symmetric cargoes, together with a ubiquitin-null control (Figure 4C). All three ubiquitinated cargoes were delivered normally in wild-type cells, so the differences in the Hse1-DUb background do not reflect a failure of ESCRT recognition per se. A remaining possibility is that cytosolic bulk sterically occludes the adjacent ubiquitin from ESCRT-0. The symmetric cargo argues against this: it carries cytosolic bulk flanking the same ubiquitin yet was incorporated efficiently.

**Figure 4.**
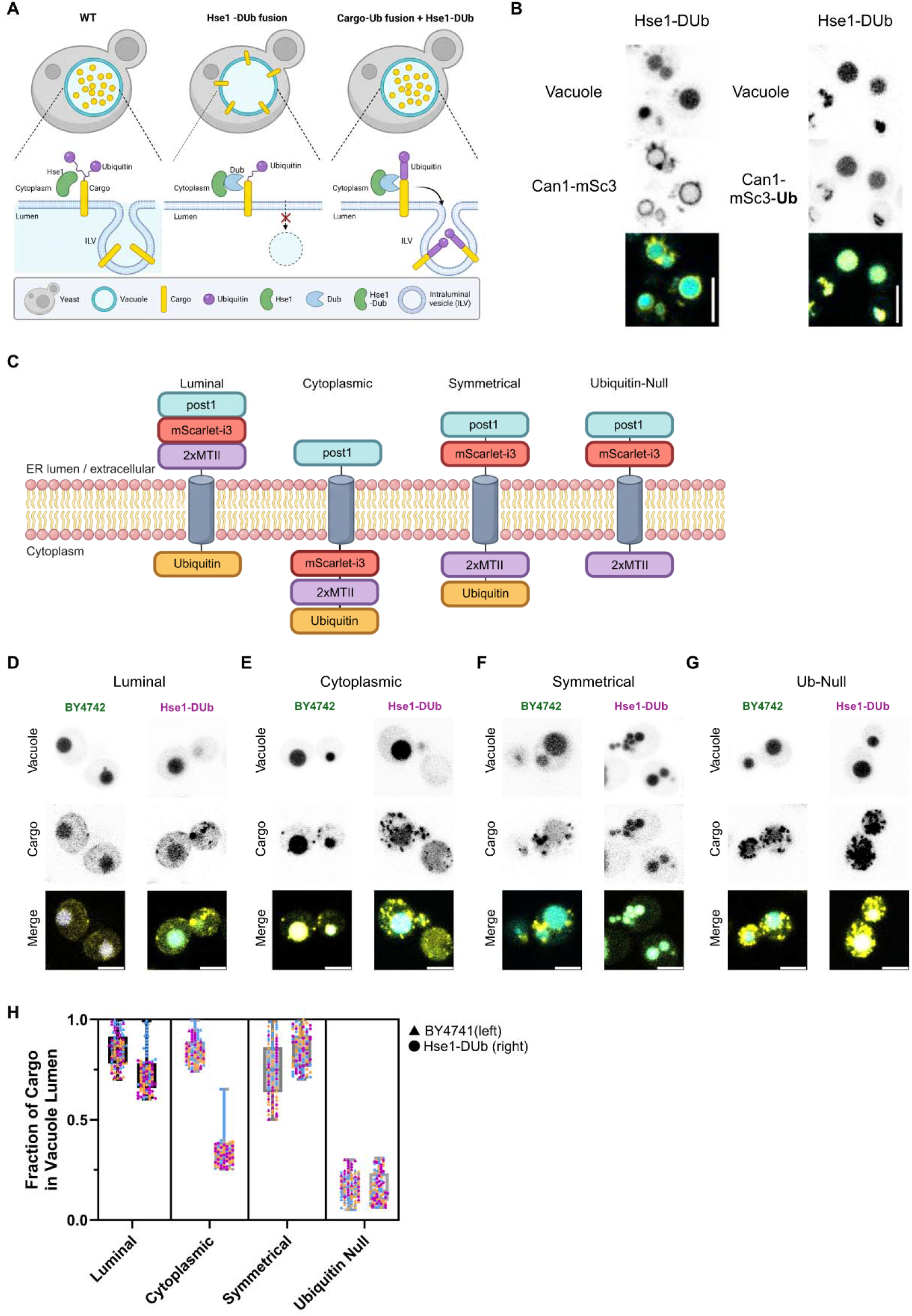
Asymmetric cargo topology affects cargo sorting efficiency in cellulo. **(A)** Schematic of the selective cargo sorting assay using Hse1p fused to a deubiquitinase (Hse1p–DUb). **(B)** Representative images showing restoration of Can1p–mScarlet3 sorting by in-frame ubiquitin fusion in the Hse1p–DUb background. **(C)** Schematic of cytoplasmic-large, luminal-large, symmetric, and ubiquitin-null cargoes, based on membrane topology predicted with Protter. **(D to G)** Representative images of luminal-large **(D),** cytoplasmic-large **(E),** symmetric **(F),** and ubiquitin-null **(G)** cargoes in BY4742 and Hse1p–DUb backgrounds. **(H)** Fraction of cargo signal in the vacuole (vacuolar/total cell signal) in BY4742 (triangles, left) and Hse1p–DUb (circles, right) backgrounds for the conditions in (D) to (G). Each symbol represents one cell (20 cells per experiment); colors denote three independent experiments. Box plots show the median (center line), interquartile range (box limits), and minimum to maximum (whiskers). Scale bars, 5 µm. Schematics created with BioRender.com.

The determining variable is therefore the distribution of bulk across the bilayer, not its presence on the cytosolic face. During ILV budding, the leaflet facing the lumen becomes the convex surface of the bud and the cytosolic leaflet becomes concave. Because crowding expands the leaflet it occupies, and an expanded leaflet preferentially occupies the convex surface, the reconstitution predicts that luminal protein bulk should favor ILV incorporation whereas cytosolic bulk should oppose it.

When expressed in wild-type cells, all three ubiquitinated cargoes—cytosolic-large, luminal-large, and symmetric—were predominantly delivered to the vacuolar lumen, whereas the ubiquitin-null cargo accumulated in cytoplasmic puncta. Thus, altering the distribution of protein bulk did not by itself prevent cargo recognition or delivery through the endosomal pathway. In the Hse1-DUb background, however, the topology of the cargo domain strongly affected ILV incorporation. Cytosolic-large cargo accumulated in endosomal puncta and showed reduced delivery to the vacuolar lumen, whereas luminal-large cargo was efficiently delivered to the vacuolar lumen. Symmetric cargo showed intermediate-to-enhanced delivery relative to the cytosolic-large construct (Figure 4D–G).

The crowding dependence of cargo incorporation is consistent with the directional response observed in the reconstituted system. When protein bulk is preferentially distributed toward the leaflet that becomes the convex surface during budding, its incorporation is favored; when bulk is concentrated on the opposing leaflet, incorporation is impaired. Because ubiquitin-dependent ESCRT recognition was maintained across the functional cargoes, these differences cannot be attributed simply to the presence or absence of the ESCRT-targeting signal. Instead, they identify the spatial distribution of cargo protein bulk across the bilayer as a variable that influences the efficiency with which cargo is incorporated into ILVs.

These observations provide a cellular correlate of the physical mechanism established by reconstitution: asymmetric protein occupancy can influence the membrane-remodeling pathway available to ESCRT-III. Thus, cargo is not only a substrate recognized by the ESCRT machinery but also contributes to the physical asymmetry of the membrane on which ESCRT-III acts.

## DISCUSSION

ESCRT-III-mediated ILV formation presents a fundamental topological problem: ESCRT-III assembles on the cytosolic surface of endosomal membranes yet drives deformation away from the cytosol. Our results indicate that this directionality emerges from the coupling between ESCRT-III polymer mechanics and membrane asymmetry. Differences in lipid composition or protein occupancy between the two leaflets alter their physical state and preferred area, generating a curvature bias that ESCRT-III polymerization can express as membrane deformation.

Bilayer asymmetry is a defining feature of cellular membranes and is absent from most reconstituted bilayers. This single difference offers a parsimonious account of the discrepancy between the orientation of ESCRT-III-mediated remodeling in cells and in vitro, without requiring any difference in the machinery itself. The direction is dictated by the difference of internal stress between the two leaflets. Our in vitro results also indicate that this stress difference can be generated by cargo crowding. In vivo, however, protein crowding and lipid asymmetry are both playing at the same time. Depending on the situation, their effects could be synergistic or antagonistic. At the plasma membrane and in the endosomes, the large extracellular domains of proteins and their glycosylation ^56,57^ and the proposed orientation of the asymmetric stress caused by lipid composition ^42,58,59^ seems to be synergistic, orienting ESCRT-III-mediated deformation towards the extracellular space and the endosomal lumen.

Our work also supports that modifications of the lipid composition may be needed to trigger membrane deformation. This is consistent with previous findings that the conversion of sphingomyelin to ceramide is needed for inward budding of ILVs ^60–62^. The phosphorylation of sphingosine - a shape conversion causing the opposite stress asymmetry in the membrane than the degradation of sphingomyelin into ceramide - is needed for endocytic budding ^63^ in a process similar to Golgi tubulation caused by PLA2 ^51^.

Overall, cells spend a considerable amount of energy to maintain membrane asymmetry ^64^ as proteins necessary to maintain the asymmetry use metabolic ATP to maintain it. Analogous to lipid gradients between organelles being used for lipid transport ^65^, this study shows that bilayer asymmetry can sort energy that can be converted to work required for membrane transport, and in particular membrane deformation.

Our reconstitutions isolate the bending step using Snf7p; how bilayer asymmetry shapes subsequent scission by the complete ESCRT-III/Vps4 machinery will be an important next step. FLIM probes resolve leaflet-specific changes in packing and mechanical state, and future approaches capable of quantifying absolute differential stress will enable a quantitative model of this coupling. In cells, where ILVs bud in a single direction, we assessed sorting efficiency as the readout of membrane asymmetry, and direct measurements of endosomal leaflet asymmetry will further refine this link. Whether this principle extends to other membrane-remodeling machines remains an open question.

## MATERIALS AND METHODS

### Reagents and resources

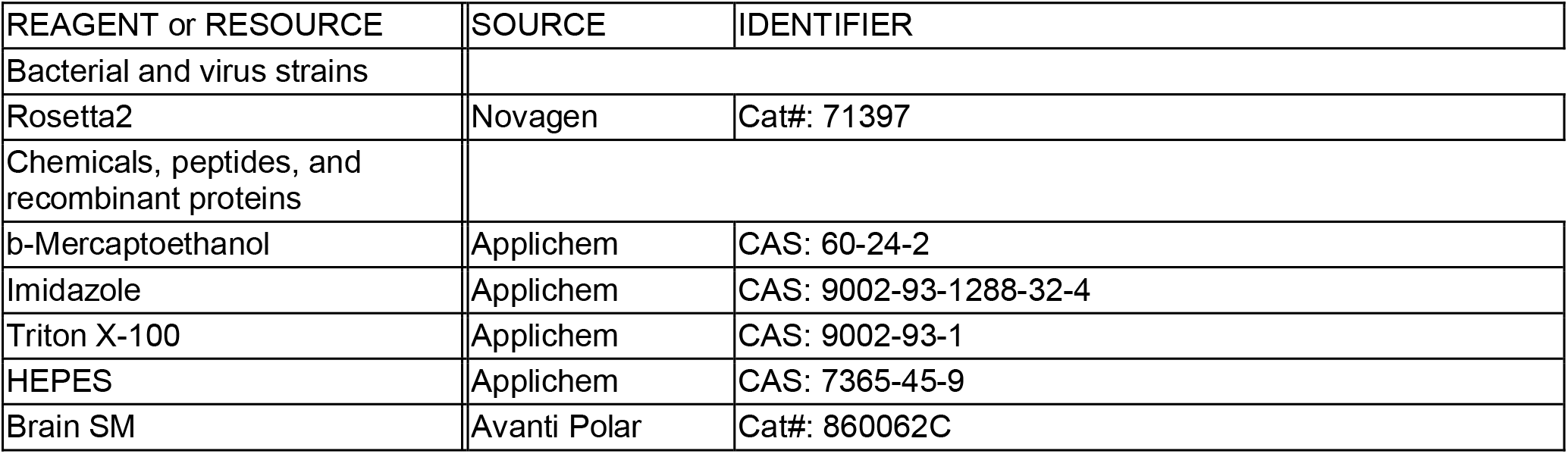
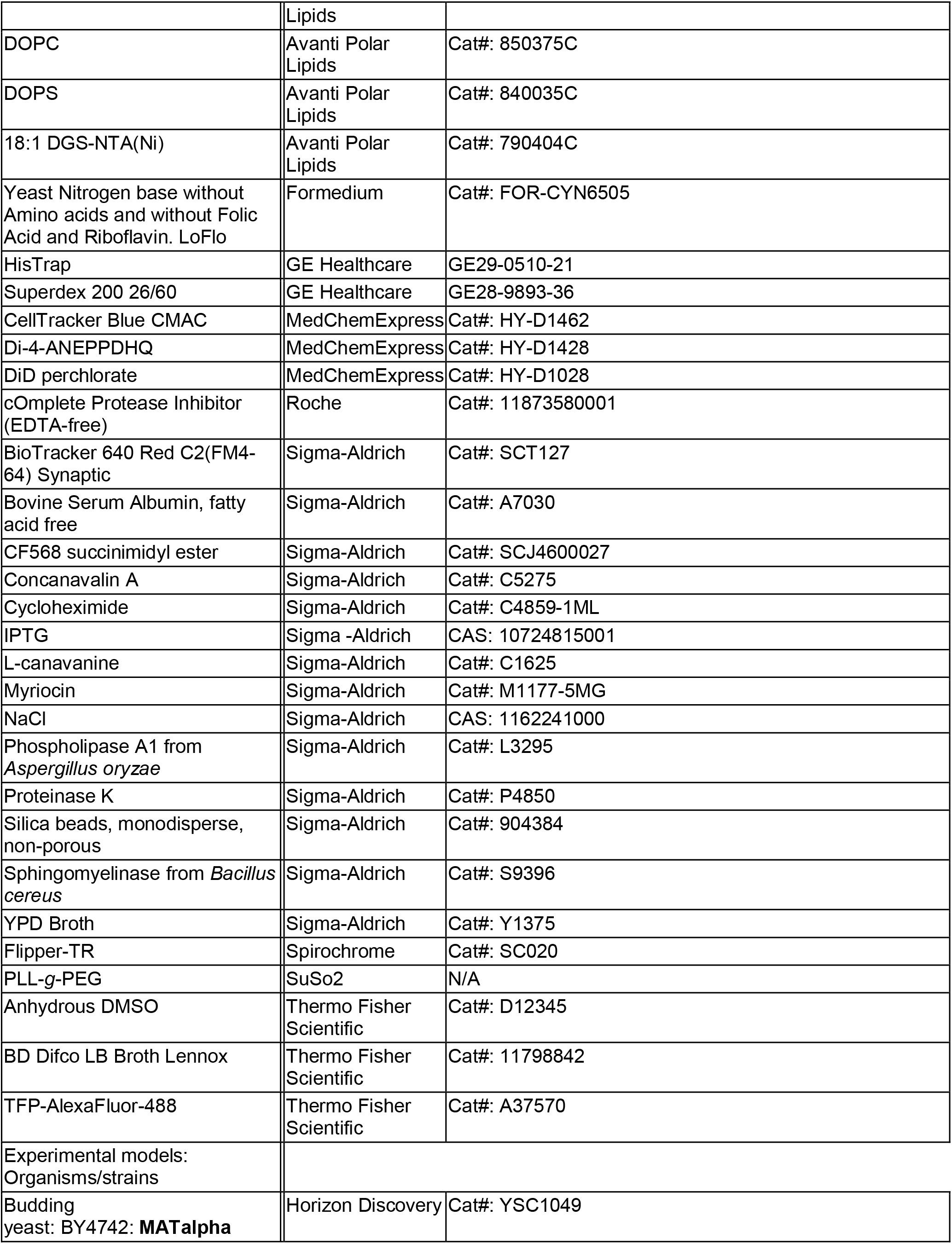
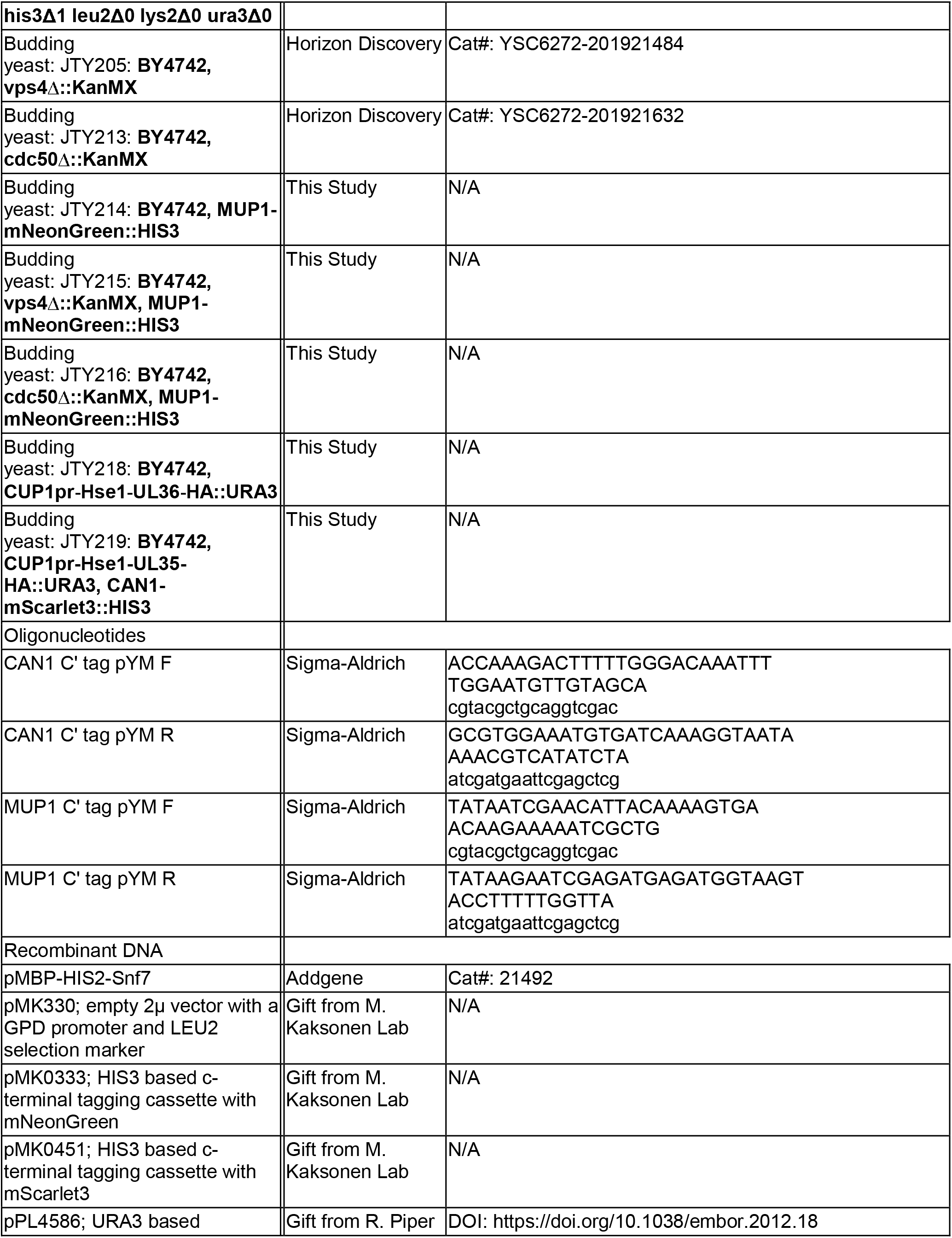
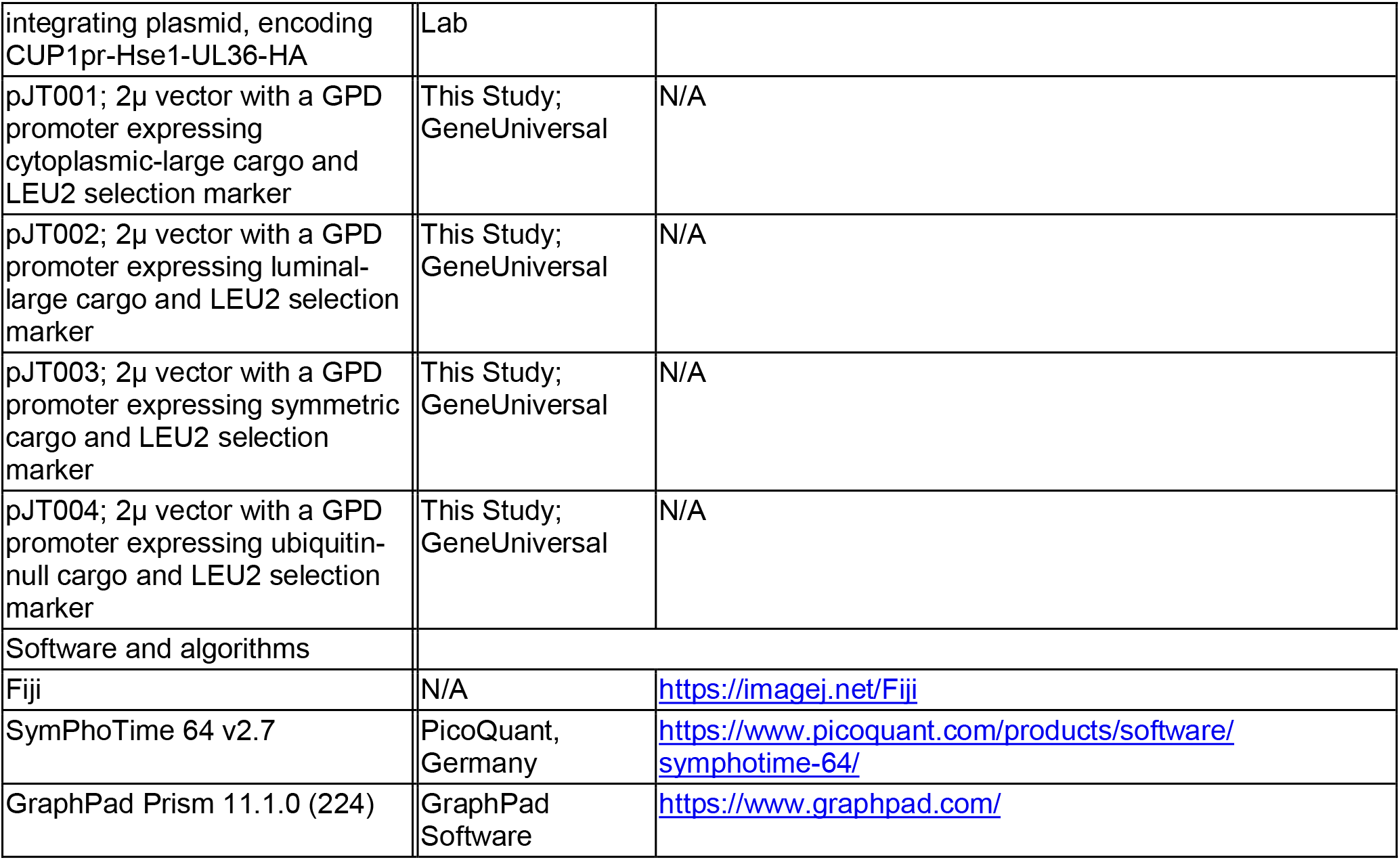

### Experimental procedures

#### Reagents

All chemical reagents were purchased from Sigma-Aldrich (St. Louis, Missouri, USA) unless otherwise indicated. Stock compounds were made in anhydrous DMSO (D12345; Thermo Fisher Scientific) unless explicitly stated. Compounds were used at the following concentrations unless explicitly stated: 600 ng/mL (2 mg/mL stock) Myriocin (M1177-5MG; Sigma Aldrich); 250 nM (1mg/mL stock) Di-4-ANEPPDHQ (HY-D1428; MedChemExpress); 1 µM (1mM stock) Flipper-TR (SC020; Spirochrome); 5.5 µM (1 mg/mL stock) BODIPY-C12 (HY-D1553; MedChemExpress); 0.05 mol% (100 µM stock) DiD perchlorate (HY-D1028; MedChemExpress); 10 µM (10 mM stock) CellTracker Blue CMAC (HY-D1462; MedChemExpress); 10 µM (10 mM stock) BioTrack 640 Red C2(FM4-64) Synaptic (SCT127; Sigma Aldrich).

All lipids were purchased from Avanti Polar Lipids. Lipid stocks were stored at −80°C in chloroform in amber glass vials and under argon gas.

Yeast Strains, Media, and Genetic Manipulation:

*Saccharomyces cerevisiae* strains used in this study are listed in the reagents and resources table. Standard LiAc/PEG-based protocols were used for genetic manipulation and transformation as previously established ^66^.

For live cell microscopy and electron microscopy, cells were cultured at 30°C (unless otherwise noted in the text) to mid-log phase in LoFlo medium (0.69% [w/v] yeast nitrogenous base w/o amino acids and without Folic Acid and Riboflavin (Formedium) and 2% [w/v] glucose) supplemented with amino acids and/or antibiotics for plasmid selection and auxotrophic complementation.

#### Plasmids

Plasmids used in this study are listed in the reagents and resources table. Protein constructs were synthesized by GeneUniversal (Delaware, USA) and subcloned into yeast expression vectors. pMBP-HIS2-Snf7 was a gift from James Hurley (Addgene plasmid # 21492 ; http://n2t.net/addgene:21492 ; RRID:Addgene_21492).

#### Cargo Sorting Assay and Live Cell Microscopy

For Mup1p sorting, cells were grown in LoFlo media (Formedium), with vehicle or chemical treatment, to mid-log phase at 30°C (unless otherwise noted in the text). To decrease signal from newly synthesized proteins, cells were treated with 100 ng/mL (100 mg/mL stock) cycloheximide (C4859-1ML; Sigma Aldrich) for 10 minutes before and during imaging. Cells were stained with CMAC and/or FM 4-64 for 30 minutes at 30°C with agitation before centrifuging at 500xg for 5 minutes to remove excess stain. Cells were washed 3 times with LoFlo medium before imaging on Concanavalin A (1 mg/mL) coated ibidi 8-well, #1.5H glass bottom chamber slides (80807; ibidi).

All yeast were imaged using a point-scanning A1 confocal microscope built on a Nikon Ti2 Eclipse inverted system (Nikon) equipped with a 100x, 1.45NA oil-immersion objective at room temperature.

#### Quantification of Vacuolar Sorting

Quantification of vacuolar protein sorting was performed in Fiji (ImageJ). Vacuoles were identified using CMAC staining of the vacuolar lumen and segmented by intensity thresholding. A whole-cell region of interest was defined for each cell, and the vacuolar region was subtracted from this mask to define the non-vacuolar cellular area.

Cargo fluorescence intensity was measured in the vacuolar lumen and in the entire cell. Vacuolar sorting efficiency was calculated as the ratio of cargo signal in the vacuolar lumen over the total cargo signal in the whole cell (including the vacuole). Background fluorescence was subtracted using regions without cells.

#### Canavanine Sensitivity and Growth Assays

Cells were pretreated with DMSO or myriocin in arginine-free synthetic defined (SD) medium for 24 h. Cells were subsequently diluted to an initial OD600 of 0.1 in 96-well plates containing arginine-free SD medium supplemented with 1 µg/mL canavanine, 600 ng/mL myriocin, or the corresponding vehicle controls, as indicated. Growth was monitored by measuring OD600 at regular intervals for 72 h at 30°C using a microplate reader.

#### Protein Purification and Labeling

Snf7 (Addgene plasmid no. 21492) was purified as previously described ^4^.

The His-Ub4 construct was subcloned into the pCoofy17 backbone to encode an N-terminal His_₆_-SUMO tag followed by a SenP1 cleavage site upstream of the linear His₆-tetraubiquitin chain as previously described ^67^. His-Ub4 was expressed in E. coli Rosetta2 (DE3) cells under kanamycin selection. Cultures were grown to OD_₆₀₀_ ≈ 0.8 and protein expression was induced with 0.5 mM IPTG, followed by overnight expression at 20 °C.

Cells were harvested and lysed by sonication in lysis buffer containing 20 mM HEPES pH 8.0, 300 mM NaCl, 30 mM imidazole, 0.5 mM TCEP, and EDTA-free complete protease inhibitors (Roche). Lysates were clarified and applied to a HisTrap affinity column. Bound protein was eluted using elution buffer (20 mM HEPES pH 8.0, 300 mM NaCl, 250 mM imidazole, 0.5 mM TCEP). Eluted fractions were dialyzed overnight at 4 °C into cleavage buffer (20 mM HEPES pH 8.0, 200 mM NaCl, 5% glycerol, 0.5 mM TCEP) to remove imidazole. SenP1 protease was added and the His_₆_-SUMO tag was cleaved overnight at 4 °C.

Cleavage products were separated by size-exclusion chromatography on a HiLoad 26/60 Superdex 200 column equilibrated in final buffer (20 mM HEPES pH 8.0, 200 mM NaCl, 5% glycerol). Fractions containing His-Ub4 were pooled and concentrated using a 30 kDa MWCO Amicon centrifugal filter.

Purified protein was fluorescently labeled using either TFP-Alexa Fluor 488 (ThermoFisher Scientific) or CF568 succinimidyl ester (Sigma-Aldrich) according to the manufacturer’s instructions.

#### Giant Unilamellar Vesicle Formation and Microscopy

GUVs were prepared by hydration of lipid films deposited on silica beads as previously described ^68–70^.

1,2-dioleoyl-sn-glycero-3-phosphocholine (DOPC) and 1,2-dioleoyl-sn-glycero-3-phospho-L-serine (sodium salt) (DOPS) were mixed at a molar ratio of 60:40 in chloroform at a total lipid concentration of 1 mM. Where indicated, additional lipids were included at the following molar percentages: DiD perchlorate (0.05 mol%) for membrane visualization, Flipper-TR and Di-4 (each 0.5 mol%) for FLIM measurements, porcine brain sphingomyelin (bSM, 5 mol%), or DGS-NTA(Ni) (11.4 mol%). In all cases, the molar fraction of DOPC was reduced accordingly to maintain a total lipid composition of 100 mol%.

Lipid mixtures were dried under a gentle stream of argon and residual solvent was removed in a vacuum oven at 30 °C for 2 h. The dried lipid film was rehydrated in 20 mM HEPES (pH 8.0) to a final concentration of 0.5 mM by vortexing for 5 min.

A 13 µL aliquot of the lipid suspension was added to 2 µL of acid-cleaned 50 µm silica beads (Sigma-Aldrich) placed on parafilm. The sample was dried to deposit the lipid film uniformly onto the bead surface.

GUVs were formed by swelling the lipid-coated beads in 10 µL of 1 M trehalose at 65 °C for 15 min, followed by immediate dilution into 200 µL of experimental buffer.

For protein encapsulation, proteins were added directly to the 1 M trehalose swelling solution at the desired concentration, and the swelling temperature was reduced to 37 °C.

To prevent unwanted binding of the GUVs to the glass, #1.5H glass coverslips were thoroughly cleaned, and bath sonicated before subjected to plasma cleaning for 15 minutes. Immediately after plasma cleaning, 0.5mg/mL of PLL-g-PEG (SuSoS) was used passivate the glass surface for a minimum of 2 hours prior to imaging. Imaging chambers were thoroughly washed 5 times with buffer prior to imaging to remove unbound PLL-g-PEG.

All GUV imaging was done on an inverted spinning-disk confocal microscope built on a Nikon C1 Digital Eclipse Modular Confocal Microscope System (Nikon) and an EM-CCD camera (Evolve, Roper Scientific). The microscope was equipped with a 100x, 1.45 NA oil-immersion objective (Nikon).

#### Confocal FLIM Imaging and Lifetime Estimation

Fluorescence lifetime imaging microscopy (FLIM) was performed as previously described ^46,47,71^. FLIM was performed using a point-scanning A1 confocal microscope built on a Nikon Ti2 Eclipse inverted system (Nikon) equipped with a 100x, 1.45NA oil-immersion objective at room temperature. Excitation utilized a 485nm diode pulse laser (PicoQuant LDH-D-C-485) and operated at 20 MHz. For Flipper-TR imaging, photons were collected continuously for 45 seconds per measurement, and emission was collected through a 600/50-nm bandpass filter. Due to the difference in brightness, for Di4 imaging, photons were collected continuously for 15 seconds, and emission was collected via the same filter as with Flipper-TR.

Lifetime decay was analyzed using SymPhoTime 64 v2.7 (PicoQuant, Germany). GUVs were manually selected from the field of view using the paint tool to exclude background. GUVs that experienced movement were excluded from the analysis. GUVs within a field of view were selected until the cumulative photon count of the measurement surpassed 10^5^ photons. Thus, individual measurements may include more than one GUV. 2-exponent reconvolution fits were done on the ROIs, adjusted to the software generated IRF, to obtain the average lifetime (τ_av intensity_). All data was plotted using GraphPad Prism, and statistical tests were done using Mann-Whitney test for significance.

#### GUV lipid remodeling and leaflet-specific Di4 labeling

GUVs were formed as described above and treated with bacterial sphingomyelinase (bSMase; 3 mU) or phospholipase A1 (PLA1; 1 mU) for 10 min. Following enzymatic treatment, Snf7-AF488 was added to the GUV suspension and allowed to polymerize for 30 min. Resulting membrane deformation was visualized by fluorescence microscopy.

For measurements of membrane properties by FLIM, Di4 was incorporated into the outer leaflet by addition to the external solution. For inner-leaflet measurements, GUVs were subjected to back-extraction of outer-leaflet Di4 using fatty-acid-free BSA (1.5% w/v final concentration) for 15 min, with three solution changes. This procedure selectively removed Di4 from the outer leaflet, leaving the probe preferentially associated with the inner leaflet. Di4 fluorescence intensity was normalized to total GUV membrane intensity, revealing an approximately 50% reduction following BSA extraction, consistent with effective removal of the outer-leaflet probe. FLIM measurements were then performed as described above.

#### Leaflet-specific cargo reconstitution and membrane deformation

GUVs containing 11.5 mol% DGS-NTA(Ni), 40 mol% DOPS, 48.45 mol% DOPC, and 0.05 mol% DiD were prepared as described above. To generate outer-leaflet-enriched cargo, fluorescent 6×His– tetraubiquitin (His-Ub4) was added externally to preformed GUVs at a final concentration of 1 µM, allowing binding to the Ni-NTA-containing outer leaflet. To generate symmetric cargo distribution, His-Ub4 was incorporated during GUV formation, allowing the construct to bind to both leaflets of the bilayer.

For inner-leaflet-restricted cargo, externally bound His-Ub4 was selectively removed by treatment with proteinase K (0.2 mg/mL final concentration) for 10 min at room temperature. Loss of external His-Ub4 was assessed by measuring the reduction in fluorescence intensity following protease treatment. The remaining His-Ub4 was therefore restricted to the inner leaflet. Membrane deformation was subsequently monitored by fluorescence microscopy, and the direction and extent of membrane tubulation were quantified as described above.

For FLIM measurements, the same cargo-loading and proteinase K treatment procedures were performed using non-fluorescently labeled His-Ub4. Di4 labeling and FLIM measurements were performed as described above.

#### Transmission Electron Microscopy of *In Vitro* Samples

LUVs were prepared by drying lipids and resuspending them in experimental buffer to a final concentration of 2.5 mg/mL. Lipids were vortexed until the solution became opaque, yielding multilamellar vesicles, which were subjected to 11 freeze–thaw cycles in liquid nitrogen to generate large unilamellar vesicles. Purified recombinant Snf7 was added to 500 nM and, where indicated, His-Ub4 was added to 500 nM. Samples were incubated for 1 h at room temperature prior to imaging.

For negative-stain electron microscopy, 4 µL of sample was applied to glow-discharged, formvar-coated 400-mesh copper grids (Electron Microscopy Services) and incubated for 1 min before blotting. Grids were washed once with isotonic HEPES buffer with sucrose to remove excess salts, followed by two washes with 10 µL drops of 2% aqueous uranyl acetate. Samples were then stained with a final 10 µL drop of uranyl acetate for 1 min, blotted, and dried prior to imaging. Imaging was performed on a Talos L120C transmission electron microscope (ThermoFisher Scientific) operating at 120 kV, equipped with an integrated CETA 16M camera.

### Statistical analysis

Data were plotted and analysed in GraphPad Prism 11.1.0 (224). In box plots, boxes show the median and interquartile range, whiskers show the minimum and maximum, and each point represents one GUV. Points are coloured by independent experimental replicate. Measurements from independent experiments were pooled for statistical analysis, with n indicating the number of GUVs and N the number of independent experiments (see figure legends for n and N per condition). Results were consistent across independent experiments for all comparisons reported. Comparisons among three or more conditions were made by ordinary one-way ANOVA with Tukey’s multiple comparisons test, and comparisons between two conditions by two-tailed Mann-Whitney test. P < 0.05 was considered significant. Exact P values are shown on the graphs; ns, not significant.

## DATA AND MATERIALS AVAILABILITY

Requests for further information and resources should be directed to the corresponding author, Aurélien Roux.

All unique/stable reagents, plasmids, and yeast strains generated in this study are available from the corresponding author with a completed materials transfer agreement.

## ACKNOWLEDGMENTS

We thank J. Espadas and J.-M. Garcia-Arcos for helpful discussions and training in model membrane and FLIM methods, respectively, and all members of the Roux lab. We thank A.-L. Boinet and M. Tettamanti for training in yeast genetics, F. Humbert and R. Caetano for assistance with protein purification, and A. Howe and S. Barrass for electron microscopy training as well as access to the Dubochet Center for Imaging (DCI), Geneva. We thank M. Doktorova and M. Deserno for helpful discussions, C. Toret for critical reading of the manuscript, the M. Kaksonen lab for tagging and expression plasmids, and the R. Piper lab for the Hse1-DUb integrating vector. A.R. acknowledges funding from the Swiss National Science Foundation (10009104, CRSII5_189996, and 310030_200793) and the European Research Council (Synergy Grant 951324-R2-TENSION).

## AUTHOR CONTRIBUTIONS

Conceptualization, J.T. and A.R.; methodology, J.T.; investigation, J.T.; visualization, J.T.; writing – original draft, J.T. and A.R.; writing – review & editing, J.T. and A.R.; funding acquisition, A.R.; project administration, A.R.; supervision, A.R.

## USE OF AI-ASSISTED TOOLS

The authors used ChatGPT (GPT-6, OpenAI; November 2026) and Claude (Claude Opus 5.5, Anthropic; November 2026) for language refinement, correction, and editing to improve clarity and concision. The authors reviewed and edited all resulting text and take full responsibility for the content of this manuscript.

## SUPPLEMENTARY INFORMATION

Figures S1–S4 (at the end of this document)

## SUPPLEMENTARY FIGURES

**Figure S1.**
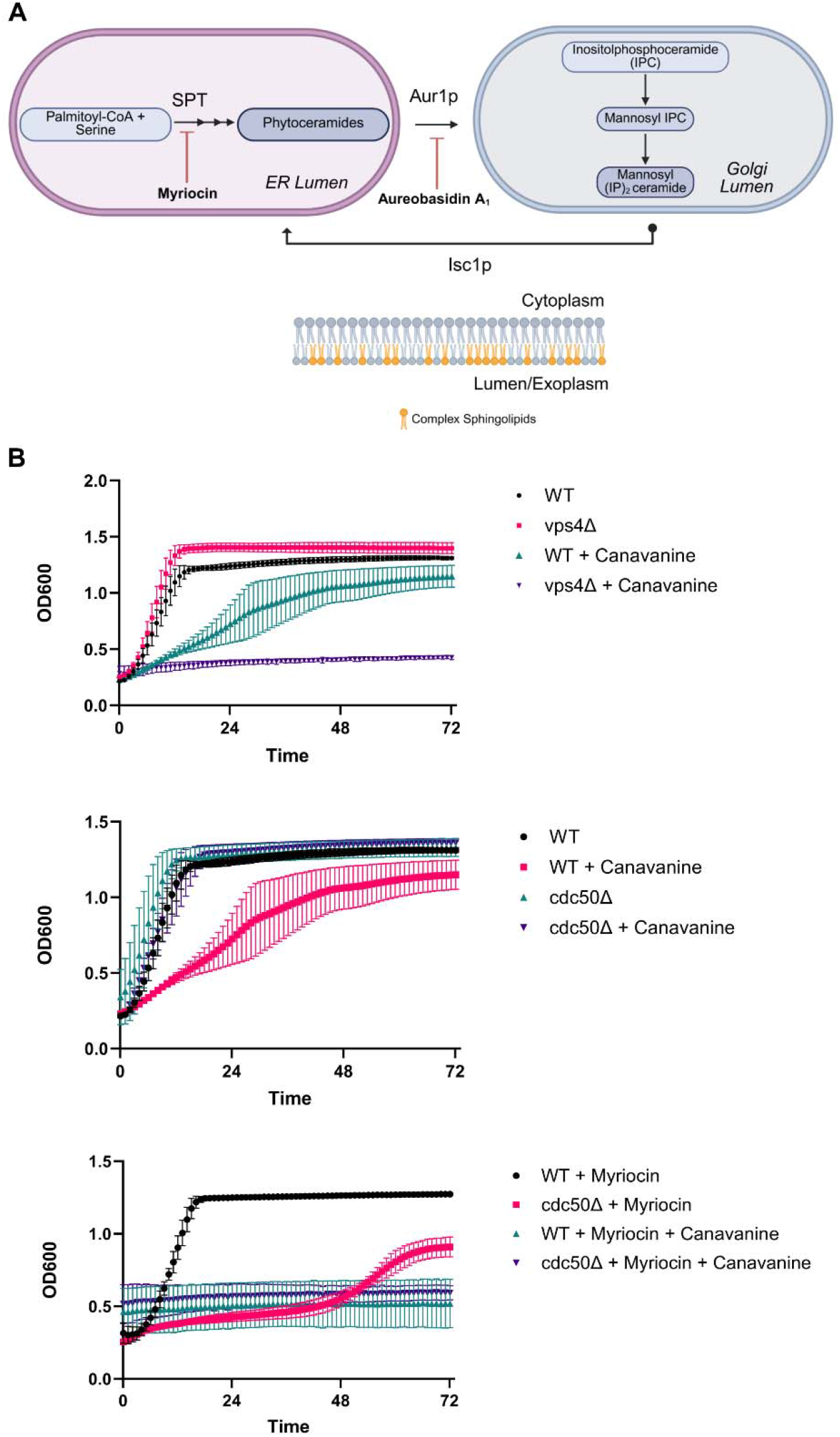
Myriocin treatment increases sensitivity to canavanine. **(A)** Schematic of the sphingolipid synthesis pathway in budding yeast and leaflet localization of sphingolipids. **(B)** Growth curves (OD_₆₀₀_ over 72 h) with or without canavanine, a toxic arginine analog taken up by the arginine permease Can1p. Top: BY4742 cells and vps4Δ mutants. Middle: BY4742 cells and cdc50Δ mutants. Bottom: BY4742 cells and cdc50Δ mutants pretreated with and grown in myriocin, an inhibitor of sphingolipid synthesis, or DMSO (vehicle). Data are mean ± SD of three biological replicates.

**Figure S2.**
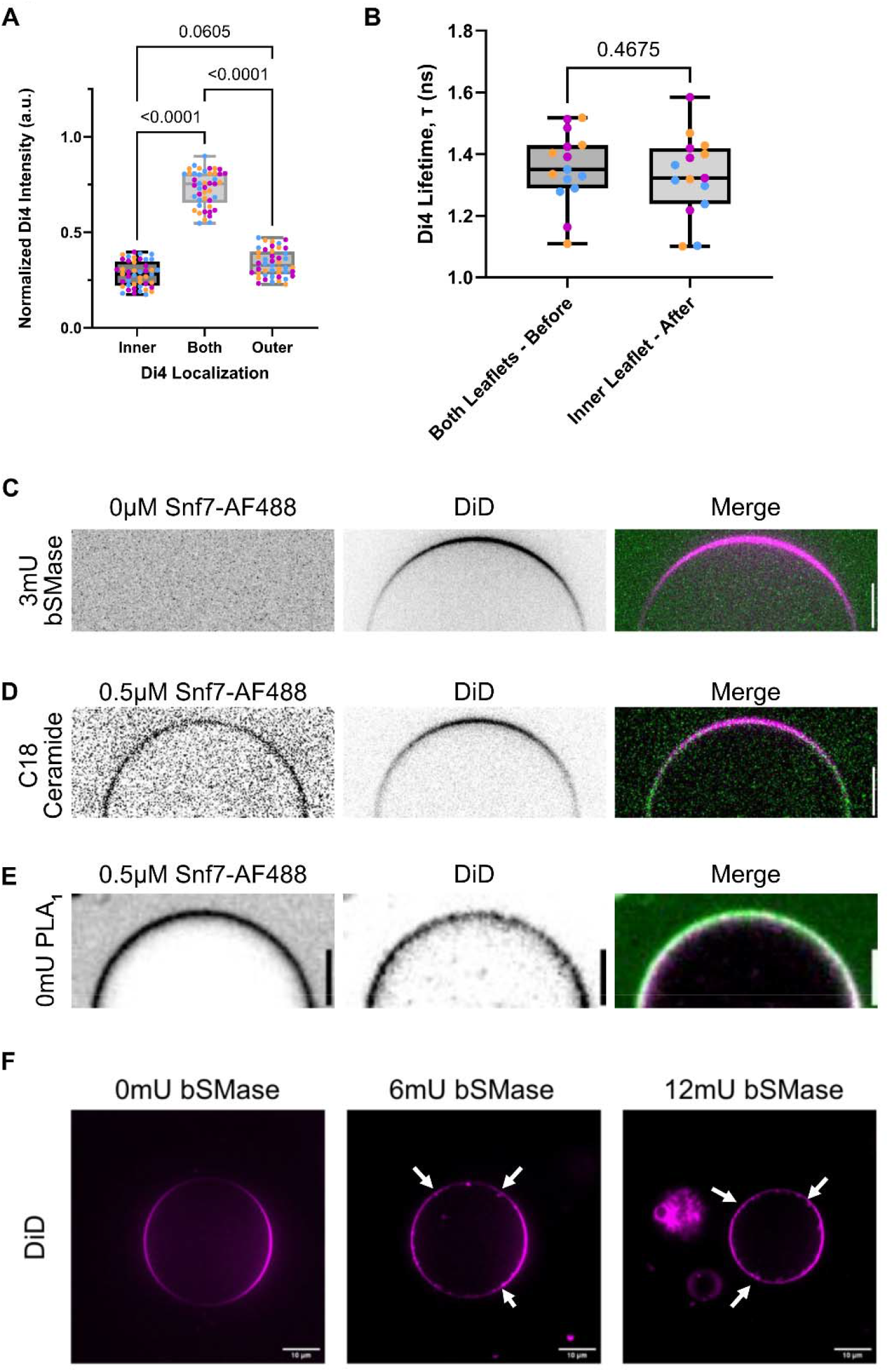
Di4 reports on leaflet-specific lipid packing, and PLA_₁_ treatment selectively alters the outer leaflet of GUVs. **(A)** Di-4 fluorescence intensity (a.u.), normalized to DiD, of GUVs with Di-4 in the inner leaflet only (Inner), both leaflets (Both), or the outer leaflet only (Outer). For Inner, Di-4 was incorporated during GUV formation and back-extracted from the outer leaflet with fatty acid–free BSA. For Both, it was incorporated during formation only. For Outer, it was added externally to preformed GUVs. **(B)** Fluorescence lifetime of Di-4 in GUVs labeled during formation, before (Both) and after (Inner) back-extraction with fatty acid–free BSA. **(C)** Representative confocal images of GUVs with Alexa Fluor 488–labeled Snf7p (0 µM), with bSMase (3 mU) in the bottom row. **(D)** Representative confocal images of GUVs containing symmetric C18 ceramide with Alexa Fluor 488–labeled Snf7p (0.5 µM). **(E)** Representative confocal images of GUVs with Alexa Fluor 488–labeled Snf7p (0 µM), with PLA1 (0 mU) in the bottom row. In (C) to (E), images show Snf7p (left), DiD (middle), and merge (right; Snf7p in green, DiD in magenta). In (A) and (B), each dot represents one GUV (10 and 5 GUVs per experiment, respectively); colors denote three independent experiments. Box plots show the median (center line), interquartile range (box limits), and minimum to maximum (whiskers). Lifetimes are in ns. Statistical significance was assessed by one-way ANOVA with Tukey’s multiple comparisons test (A) or two-sided Mann–Whitney U test (B); ns, not significant. Scale bars, 5 µm. **(F)** Representative confocal images of GUVs treated with 0mU, 6mU, and 12mU of bacterial sphingomyelinase. Scale bars, 10 µm.

**Figure S3.**
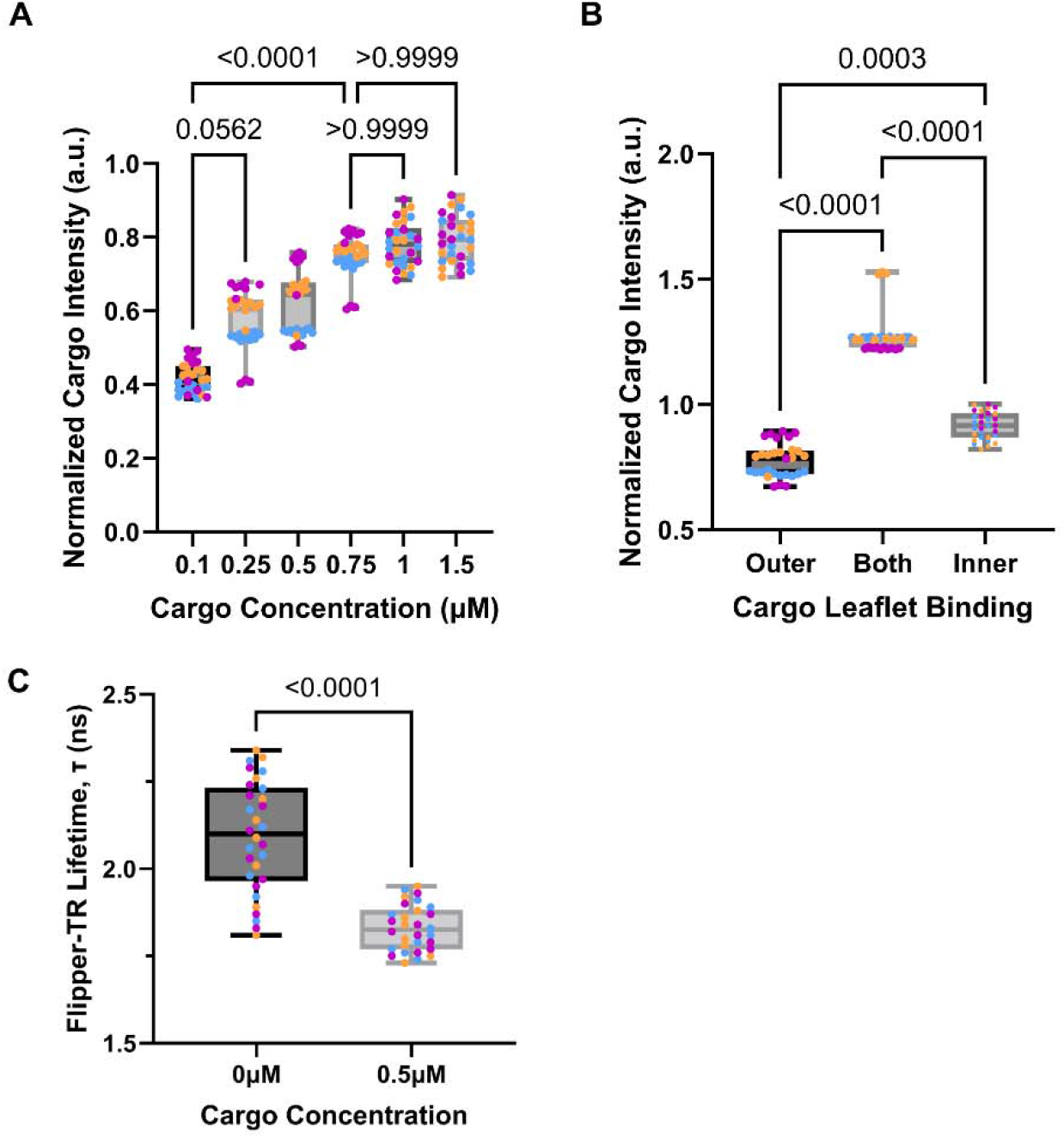
Characterization of His-Ub4 cargo binding to asymmetric GUVs. **(A)** Binding of His-Ub4 cargo to GUVs at 0.1 to 1.5 µM, with cargo fluorescence intensity normalized to DiD. **(B)** Normalized cargo intensity with His-Ub4 added externally to preformed GUVs (Outer), incorporated into both leaflets during GUV formation (Both), or incorporated into both leaflets and digested with proteinase K to remove the outer-leaflet pool (Inner). **(C)** Fluorescence lifetime of Flipper-TR in GUVs with 0 or 0.5 µM non-fluorescent His-Ub4 bound to the outer leaflet only. Each dot represents one GUV (10 GUVs per experiment); colors denote three independent experiments. Box plots show the median (center line), interquartile range (box limits), and minimum to maximum (whiskers). Lifetimes are in ns. Statistical significance was assessed by one-way ANOVA with Tukey’s multiple comparisons test (A and B) or two-sided Mann–Whitney U test (C) (n = 30 GUVs per condition); P values are shown on the plots.

**Figure S4.**
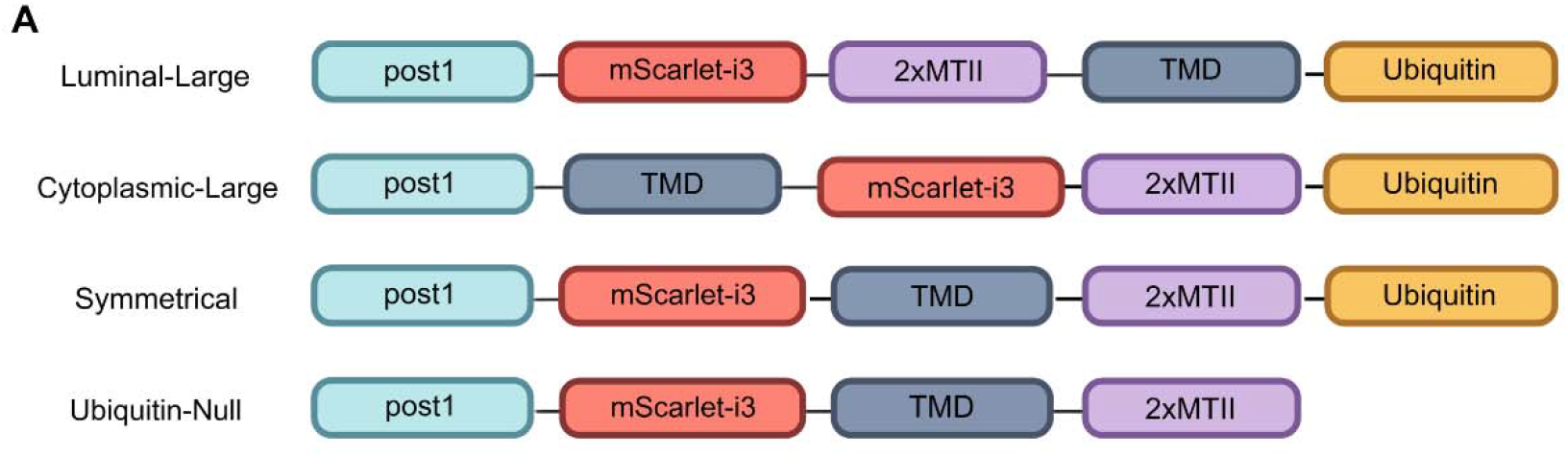
Design and predicted membrane topology of synthetic ESCRT cargoes. **(A)** Domain organization of the four synthetic cargoes: Luminal-Large, Cytoplasmic-Large, Symmetrical, and Ubiquitin-Null. Each construct contains pOst1 signal sequence, the fluorescent protein mScarlet-i3, two copies of mouse metallothionein II (MTII), and the transmembrane domain (TMD) of Cps1p. Ubiquitin is fused at the C-terminus of every construct except Ubiquitin-Null. Domains are shown in order from N- to C-terminus. Colors: pOst1 (cyan), mScarlet-i3 (red), 2xMTII (purple), Cps1p TMD (gray), ubiquitin (orange). Schematic made in BioRender.

